# Demographic variability in spruce-fir-beech forest stands in Europe

**DOI:** 10.1101/120675

**Authors:** Guillaume Lagarrigues, Franck Jabot, Andreas Zingg, Jean-Claude Gégout, Matija Klopčič, Benoît Courbaud

## Abstract

Many studies have predicted large changes in forest dynamics during the next century because of global warming. Although empirical approaches and studies based on species distribution models provide valuable information about future changes, they do not take into account biotic interactions and stand-level demographic variations. The objective of this study was to quantify the local and regional variability of the growth and regeneration of three important forest species growing often in mixed stands in Europe (*Picea abies (L.) Karst., Abies alba Mill., Fagus sylvatica*), and to assess the climatic drivers of this variability. For that purpose, we collected a large forestry data set compiling the long-term (up to 100 years) evolution of species and size distributions for 163 stands across Europe, in the mesic distribution area of these forests. We used an inverse modeling approach, Approximate Bayesian Computation, to calibrate an individual-based model of forest dynamics on these data. Our study revealed that the variability of the demographic processes was of the same order of magnitude between stands of a same forest as between different forests. Out of the three species and two demographic processes studied, only the fir growth strongly varied with temperature. Water availability did not explain any demographic variation over stands. For these forests experiencing mesic conditions, local unmeasured factors seem therefore to have an influence at least as important as macro-environmental factors on demographic variations. Efforts to include these important factors in projection scenarios should therefore be prioritized. Besides, our study demonstrates that inverse modelling methods make possible the analysis of long-term forestry data. Such data should therefore be more widely compiled and used for ecological and global change research.

## 3 Introduction

The effects of current global warming on forest dynamics are currently intensively studied (Keenan 2015). Several empirical approaches are used to study the relations between climate and tree demography in order to improve projections of future forest composition. For instance, dendroecological studies use tree core samples to assess the sensitivity of annual tree growth of different species to climatic factors (Rolland et al. 1998; Carrer et al. 2012). Another example is the use of presence/absence data derived from large scale inventories to calibrate species distribution models (SDMs), which are then used to carry out projections of species distribution ranges under future climate (Piedallu et al. 2009). These two types of analysis have provided valuable information about future forest changes but both suffer from the same important drawbacks: they give little information about key population processes such as natural regeneration (Lenoir et al. 2009) and biotic interactions (Wisz et al. 2013).

In that context, dynamic vegetation models (DVMs) that accurately represent the demographic processes in tree populations are now recognized as essential tools to account for the complexity of forest dynamics (Fontes et al. 2010; Hickler et al. 2012). They are driven by competition between individuals or species, so that predicted dynamics depends on the species composition, size structure and tree density of forest stands. These models have experienced significant developments during the last decades, with a better consideration of climatic drivers (Bugmann and Solomon 2000), couplings with SDMs (Thuiller et al. 2014) or integration of multi-scale processes, such as forest landscape models (Wang et al. 2013) or second generation dynamic global vegetation models (Scheiter et al. 2013). These process-based models have provided new knowledge about population dynamics and are now used to predict species range shifts (Snell et al. 2014) and future species composition of forests (Elkin et al. 2015). However, the species responses to future climate given by DVMs are still highly uncertain (Lindner et al. 2014; Tegel et al. 2014), mainly because of inaccurate model calibration. The parametrization of these models has been generally carried out by calibrating each process independently, using specific data sets or published literature (Bugmann 1996; Snell et al. 2014; Courbaud et al. 2015). Inverse calibration techniques such as Approximate Bayesian Computation (ABC) offer new opportunities to improve the calibration of DVMs. Such flexible techniques enable to use a wider range of data to calibrate a model than classical statistical approaches: a complex DVM can thus be calibrated using population-level data. A second advantage of the inverse calibration techniques is that they enable to calibrate jointly the different demographic processes and therefore to improve model projections by taking into account possible interactions between various demographic processes (Hartig et al. 2012; Lagarrigues et al. 2015).

In this study, we focused on uneven-aged mountain forests dominated by Norway spruce (*Picea abies (L.) Karst*.), Silver fir (*Abies alba Mill.*) and European beech (*Fagus sylvatica*), the dominant tree species in European mountain ranges. Uneven-aged forest stands, where trees of all ages coexist, are interesting study cases for demographic analyzes because the different demographic processes occur continuously over long time periods in these stands. Spruce-fir-beech forests are naturally present or have been planted in Western Europe from 600 m up to 2000 m above sea level. Given the broad distribution of these forests, the influence of climate on the demography of spruce, fir and beech has been intensively studied (Mäkinen et al. 2002; Dittmar et al. 2003; Csillery et al. 2013; Körner et al. 2016). If many dendroclimatological or productivity studies have shown that the growth of these species is generally more limited by temperature than by water availability (Wilson and Hopfmueller 2001; Dittmar et al. 2003; Büntgen et al. 2007; Seynave et al. 2008), others have highlighted their high sensitivity to drought (Pinto et al. 2008; Battipaglia et al. 2009; Carrer et al. 2012). Consequently, some SDMs have projected that the distribution areas of these species could be strongly reduced by year 2100 because of climate change (Piedallu et al. 2009), to the benefit of more drought-tolerant species. Yet, some studies questioned the sensitivity of spruce, fir and beech to a warmer and drier climate (Tinner et al. 2013; Tegel et al. 2014). Moreover, the renewal of uneven-aged forests is based on natural regeneration, and the relations between this tree life stage and climate are still poorly known (Lenoir et al. 2009). Indeed, the conditions favorable to seed production may be different than those favorable to seed germination or sapling growth and survival. Although some studies managed to assess climate effects on seed production (Mencuccini et al. 1995; Selås et al. 2002), the quantification of these effects on regeneration processes remain difficult because of the simultaneous influence of other environmental factors, such as the presence of micro-sites for seed germination (Hunziker and Brang 2005; Paluch and Jastrzębski 2013) or the ground light conditions for sapling growth and survival (Grassi et al. 2004; Stancioiu and O’Hara 2006).

DVMs commonly used to study European mountain forests, such as ForClim (Bugmann 1996) or PICUS (Lexer and Honninger 2001; Seidl et al. 2005), represent the relationship between tree diameter growth and temperature by a parabolic increasing response is currently used to link tree diameter growth to temperature, with an asymptote for high temperatures in ForClim (Bugmann and Solomon 2000). The response of tree recruitment flow to climate is either modeled with functional forms similar to tree growth (Lexer and Honninger 2001) or by rectangular responses (*i.e* a constant effect between two thresholds and no regeneration outside that range) (Bugmann 1996), with an additional threshold effect of winter temperature, in order to consider the sensitivity of saplings to extreme cold conditions. These theoretical responses lead to satisfactory predictions in terms of species occurrences and abundances (Bugmann and Solomon 2000; Seidl et al. 2005). They have however failed to explain some aspects of forest dynamics such as successional patterns (Didion et al. 2009). DVMs have predicted significant changes in forest dynamics under climate change (Elkin et al. 2013). Assessing the robustness of these predictions is therefore of major environmental and economic importance (Hanewinkel et al. 2013). In particular, it is necessary to clarify whether the observed variability of species dynamics within their distribution areas can be explained by climatic drivers or whether other drivers possibly interacting with climate change have been overlooked (Carcaillet and Muller 2005; Granier et al. 2007; Didion et al. 2011; Bodin et al. 2013).

The goal of this study was to document the contribution of climate to the variability of the forest dynamics in uneven-aged European spruce-fir-beech mountain forests. For that purpose, we used a set of long-term forestry data collected in a large network of 163 stands located in 19 forests representing a large gradient of climatic conditions within the distribution area of the three species (*P. abies, A. alba* and *F. sylvatica*). From these data, we assessed the stand dynamics using two methods: first, we computed population level demographic fluxes directly from the data; second, we calibrated the individual-based forest DVM Samsara2 (Courbaud et al. 2015), using the inverse modeling technique developed by Lagarrigues *et al*. (2015). This approach enabled us to precisely identify the effect of climate on each demographic process at the individual tree level and to factor out the multiple stand level endogenous factors (such as tree density, species composition or size structure) that affect stand dynamics and that may blur climate impacts. We expected that: (1) tree-level processes (tree growth and recruitment) that are inferred by the model-based approach would be more strongly influenced by environmental factors than raw population-level statistics; (2) tree growth would be positively correlated with temperature and only poorly correlated with water availability in these mesic mountain forests; (3) the response of tree recruitment to macro-environmental variables would show either the same trends as tree growth, or no trend because of the prevailing role of other factors, such as micro-site availability or micro-topography.

## 4 Materials and methods

### 4.1 Study forests and forestry data

A large number of managed forests distributed across a wide geographical area were considered (Fig. 1). Our data were collected in forest management archives. In France, the French Forest National Office (ONF) manages a large number of public mountain forests and was therefore our main data source. Three other data sources were used in this study: a private forest management group located in the Jura mountains; the Swiss Federal Institute for Forest, Snow and Landscape Research (WSL) at Birmensdorf, Switzerland, which provided monitoring data collected in 16 permanent plots (Zingg et al. 2009); and the archives of Slovenia Forest Service from which data for the Slovenian forests were derived by the University of Ljubljana (partly described by Klopcic *et al*. (2010)).

**Figure 1:**
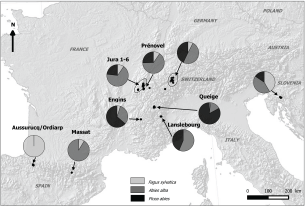
Forest locations and species composition. The pie charts represent the species relative abundances. For the sake of clarity, some forests have not been labeled, see table 1 for full information about the study forests.

These forestry data consist in two main elements: (1) forest inventories, in which information about tree abundance and species composition is given; (2) harvest reports, guvung information about the quantity and the size of trees harvested in the forest, usually on an annual basis. The merchantable volume is the usual unit of measurement used to quantify the tree size in these data. We therefore used this unit throughout this study and referred to it simply by “volume”. All recorded forests were subjected to an uneven-aged management during the data period and during several decades before the oldest inventory. We assumed that all dead trees were harvested and recorded in harvest reports. In the forest inventories, only trees above a diameter at breast height (DBH) threshold were monitored (7.5, 10 and 17.5 cm for Swiss, Slovenian and French forests respectively). To avoid a “demographic gap” when reproducing the forest history, the young stages (*i.e*. trees and saplings under the DBH threshold) in the initial states were simulated using Samsara2 (further information is given in the Appendix C). Finally, as there is no spatial information in the data, the tree positions were randomly drawn before each simulation. The data imputation procedures concerning spatial positions and young stages have been shown to only have moderate effects on simulation outcomes (Courbaud et al. 2015).

Initially, data were available for 274 stands (Appendix A, Table A.1), which are subdivisions of the 23 study forests corresponding to the management units defined by forest managers. Several filters were then applied to reach the scope of this study. First, as we restricted the study to spruce-fir-beech forests, we kept only stands where the cumulated volume of these three species was greater than 80% of the complete stand volume. In these stands, the species other than spruce, fir and beech (hereafter simply called “other species”) were then ignored, *i.e*. we removed all trees from other species from inventory and harvest data and considered that the stands contained exclusively spruce, fir and beech trees. By limiting our selection to the stands where no more than 20% of the total volume was provided by other species, we assumed that the impact of the artificial reduction of tree density by the removal of trees of other species was negligible. Second, we selected only the stands with at least 30 years between the first and the last inventory. Finally, after preliminary analysis (not detailed in this paper), we excluded the stands with aberrant data (*e.g*. unrealistic low or high numbers of trees reported in statistical inventories). In the end, we selected 163 stands in 19 forests (Table 1).

**Table 1:** Data summary. Each line is a site, which grouped various *numbers of stands* (see the Material and methods section and the Appendix A for more information about site definition). *Data period* is the mean time between the oldest and the youngest inventories. Annual volume increment and annual recruitment rate are computed on trees whose DBH > 17.5 cm (> 20 cm for Slovenian forest) and are formatted as “mean (min / max)”. The abbreviations S, F, B and O for the *species abundances* mean respectively Spruce, Fir, Beech and Other species; the abundances are given in % of the total volume. SGDD = annual Sum of Growing Degree-Days; CWB = Climatic Water Budget (see the main text for computation details).

| Country | Forest name | Site name | Site area (ha) | Number of stands | Data period (years) | Annual volume increment (m<sup>3</sup>.ha<sup>-1</sup>.year<sup>-1</sup>) | Annual recruitment rate (nb.ha<sup>-1</sup>.year<sup>-1</sup>) | Species proportions (S / F / B / O) | Elevation (m) | SGDD (°C.days) | CWB (mm) |
| --- | --- | --- | --- | --- | --- | --- | --- | --- | --- | --- | --- |
| France | Queige | Queige 1 | 19 | 1 | 49 | 4.4 (4.4 / 4.4) | 0.4 (0.4 / 0.4) | 85 / 8 / 7 / 0 | 939 | 1582 | 754 |
|  |  | Queige 3 | 224 | 6 | 49 | 4.9 (3.8 / 5.6) | 0.5 (-0.1 / 1.5) | 73 / 26 / 1 / 0 | 1294 | 1160 | 786 |
|  |  | Queige 4 | 172 | 8 | 58 | 4.3 (2.9 / 5.6) | -0.7 (-3.4 / 1.5) | 84 / 16 / 0 / 0 | 1497 | 985 | 833 |
|  |  | Queige 5 | 134 | 4 | 58 | 3.3 (2.1 / 4.5) | -0.3 (-0.4 / -0.1) | 98 / 2 / 0 / 0 | 1660 | 829 | 955 |
|  |  | Queige 7 | 33 | 1 | 83 | 4.4 (4.4 / 4.4) | 1.4 (1.4 / 1.4) | 70 / 15 / 14 / 0 | 1364 | 1208 | 757 |
|  |  | Queige 9 | 37 | 1 | 83 | 3.9 (3.9 / 3.9) | 0.3 (0.3 / 0.3) | 85 / 15 / 0 / 0 | 1651 | 914 | 822 |
|  | Lans-le-Bourg | Lans 1 | 180 | 6 | 58 | 4.3 (2.5 / 6.9) | 0.7 (-2.1 / 3.3) | 41 / 52 / 0 / 7 | 1613 | 906 | 360 |
|  | Engins | Engins 1 | 35 | 3 | 73 | 3.2 (2.6 / 3.9) | 5.9 (1.9 / 11.3) | 63 / 31 / 5 / 0 | 1312 | 1186 | 450 |
|  | Jura | Jura 1 | 168 | 19 | 34 | 8.7 (3.7 / 11.2) | 1.9 (-0.7 / 5.2) | 3 / 82 / 10 / 6 | 536 | 1948 | 625 |
|  |  | Jura 2 | 67 | 10 | 33 | 4.8 (3.2 / 6.1) | 5.3 (3.3 / 7.7) | 11 / 72 / 10 / 7 | 600 | 1782 | 778 |
|  |  | Jura 3 | 73 | 7 | 34 | 8.4 (6.2 / 10.9) | 2.8 (1.4 / 3.7) | 62 / 31 / 4 / 4 | 661 | 1700 | 793 |
|  |  | Jura 4 | 51 | 2 | 30 | 8.2 (7.5 / 8.9) | 3.3 (3 / 3.5) | 38 / 52 / 2 / 8 | 600 | 1864 | 907 |
|  |  | Jura 5 | 108 | 5 | 36 | 6.5 (4.5 / 10.5) | 2.3 (0.8 / 3.6) | 43 / 26 / 18 / 14 | 552 | 1861 | 707 |
|  |  | Jura 6 | 21 | 2 | 30 | 8.8 (7.9 / 9.7) | 6.4 (2.8 / 9.9) | 40 / 53 / 4 / 2 | 546 | 1870 | 711 |
|  | Prénoval | Prénoval 1 | 384 | 34 | 58 | 7.9 (5.6 / 12.2) | 4.1 (2.3 / 7.3) | 25 / 65 / 10 / 0 | 957 | 1419 | 1041 |
|  | Aussurucq | Aussurucq 1 | 215 | 11 | 48 | 4.3 (1.6 / 8.9) | 5.3 (1.7 / 12.6) | 0 / 0 / 96 / 4 | 623 | 2192 | 623 |
|  |  | Aussurucq 2 | 194 | 7 | 48 | 3.2 (1.3 / 6.7) | 3.8 (0.5 / 9.3) | 0 / 0 / 97 / 3 | 807 | 1918 | 631 |
|  | Massat | Massat 1 | 37 | 3 | 39 | 3 (1.4 / 4.4) | 1.5 (0.7 / 3) | 0 / 90 / 8 / 1 | 1149 | 1728 | 530 |
|  | Ordiarp | Ordiarp 1 | 174 | 6 | 50 | 3.4 (2.4 / 4.3) | 2.8 (1.5 / 3.9) | 0 / 0 / 93 / 7 | 570 | 2270 | 600 |
|  |  | Ordiarp 2 | 160 | 6 | 48 | 4 (1.6 / 6.6) | 4.6 (1.8 / 8.1) | 0 / 0 / 100 / 0 | 828 | 1927 | 700 |
| Slovenia | Slovenia | Slovenia 1 | 37 | 2 | 45 | 9.6 (6.8 / 12.4) | 9.1 (5.8 / 12.3) | 16 / 73 / 9 / 2 | 872 | 1336 | 630 |
|  |  | Slovenia 2 | 37 | 3 | 47 | 6.3 (6 / 6.8) | 10.5 (5.4 / 16.8) | 14 / 14 / 66 / 6 | 1275 | 877 | 783 |
| Switzerland | Landiswil | Landiswil 1 | 7 | 1 | 95 | 8.3 (8.3 / 8.3) | 11.7 (11.7 / 11.7) | 37 / 58 / 4 / 1 | 930 | 1452 | 320 |
|  | Lauperswil | Lauperswil 1 | 2 | 2 | 97 | 8.9 (8.6 / 9.3) | 6.3 (6.1 / 6.4) | 32 / 42 / 23 / 3 | 890 | 1427 | 239 |
|  | Niederhünigen | Niederhünigen 1 | 1 | 5 | 101 | 7.8 (6.1 / 8.7) | 8.3 (3.4 / 11.4) | 41 / 52 / 5 / 2 | 957 | 1400 | 379 |
|  |  | Niederhünigen 2 | 3 | 1 | 91 | 8.7 (8.7 / 8.7) | 9.2 (9.2 / 9.2) | 19 / 79 / 2 / 0 | 570 | 1764 | 292 |
|  | Oberlangenegg | Oberlangenegg 1 | 2 | 1 | 81 | 10 (10 / 10) | 7.8 (7.8 / 7.8) | 45 / 50 / 4 / 0 | 920 | 1409 | 425 |
|  | Röthenbach | Röthenbach 1 | 2 | 1 | 80 | 8.8 (8.8 / 8.8) | 9 (9 / 9) | 9 / 74 / 17 / 0 | 1068 | 1262 | 448 |
|  | Rougemont | Rougemont 1 | 4 | 2 | 85 | 6.9 (6.7 / 7) | 6.3 (5.7 / 6.9) | 50 / 47 / 1 / 2 | 1249 | 1280 | 1081 |
|  | Sigriswil | Sigriswil 1 | 2 | 2 | 83 | 3.8 (3.6 / 4) | 2.4 (2.2 / 2.7) | 100 / 0 / 0 / 0 | 1388 | 940 | 604 |
|  | Chenit | Chenit 1 | 2 | 1 | 79 | 3.2 (3.2 / 3.2) | 4.7 (4.7 / 4.7) | 88 / 5 / 2 / 5 | 1345 | 978 | 1303 |

### 4.2 Macro-environmental data

Monthly temperature and rainfall data were used to compute climatic indices. These data were extracted from the METEO FRANCE SAFRAN model (Durand et al. 1993) for the French and Swiss stands, and from the WorldClim data (Hijmans et al. 2005) for the Slovenian stands. The SAFRAN data were available annually from 1959 to 2013. These meteorological data were averaged for each forest stand over the time period that was common to the SAFRAN data and to the forestry stand data (*e.g*. for the Queige stands, the forestry data period was generally 1931-1980, the meteorological data were then averaged over the period 1959-1980). Worldclim climatic variables were averaged over the 1960-90 period. Because the meteorological data were of a too coarse spatial resolution, we corrected the monthly temperatures by the effect of elevation using linear regressions applied on climatic data subsets (moving window techniques used by Kunstler *et al*. (2011); mean correction = -0.0051 K.m^-1^, see Appendix B in Supporting Information for further details on this correction).

The thermal conditions of the stands were then described by an index broadly used in the literature to analyze forest dynamics: the annual sum of growing degree-days (SGDD), computed as the sum of the degrees over 5.56°C over all year, computed from mean monthly temperatures (Kunstler et al. 2011).

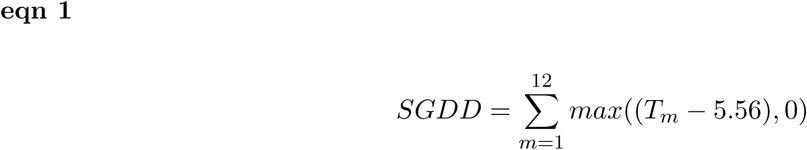
where *m* is the month index and *T_m_* the average temperature during the month *m*.

The water conditions were described by the climatic water budget (CWB) index, computed as the difference between annual total rainfall and annual potential evapotranspiration (PET). CWB was identified as a good and efficient moisture index when water availability does not limit forest dynamics (Piedallu et al. 2013), which is the case for the forests in this study (Fig. 2). The PET values were computed at the stand scale for each month from temperature and latitude data, using the formula of Oudin (Oudin et al. 2005).

**Figure 2:**
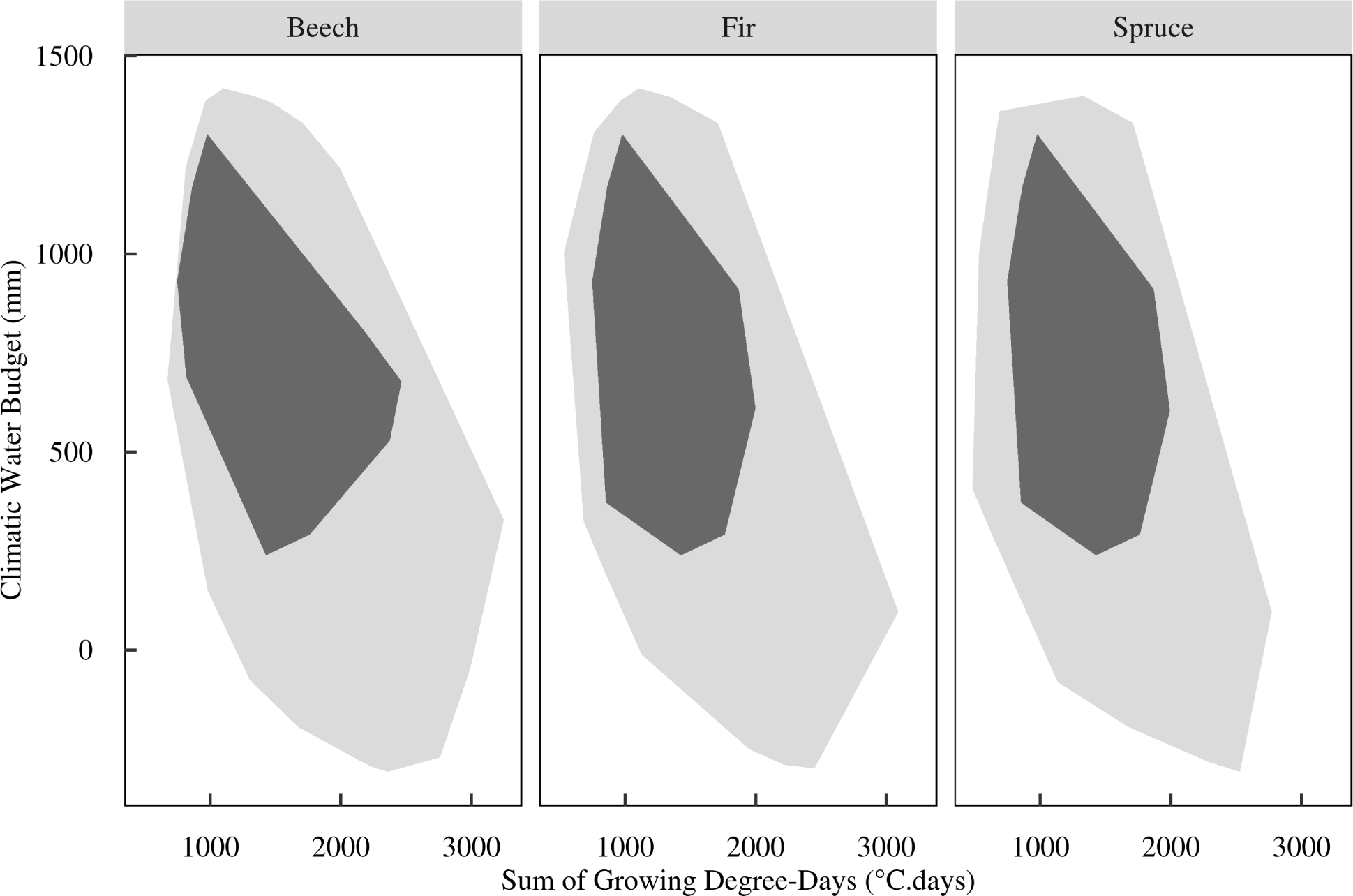
Climatic envelopes for the study species. The gray areas represent the climatic conditions where the species are present in France (based on the French National Inventory). The black areas represent the climatic conditions where the species are present in the study stands, including the forests that are outside France.

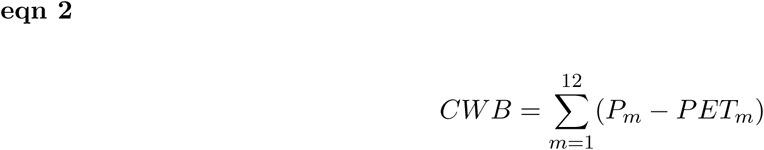

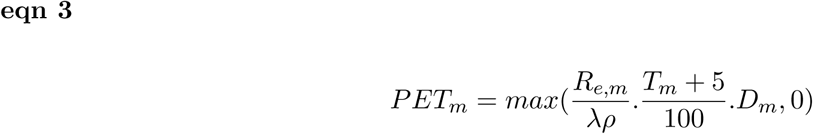
where *P_m_* is the monthly sum of precipitation, *R_e,m_* is the extraterrestrial radiation (depending on month m), is the latent heat flux (taken equal to 2.45 MJ.kg^-1^), is the density of water (1000 kg.m^-3^) and *D_m_* is the number of days in the month *m*. For further information about the computation of the different elements of this formula, please refer to Oudin *et al*. (2005) and Allen *et al*. (1998).

This simplistic PET formula may be questioned but, in the climatic area of our study, the Oudin PET values were highly correlated with the PET values computed with the formula of Penman (Penman 1948), which is considered as the best approximation of PET. We used this formula because it allowed to correct PET as a function of elevation (using the temperature correction mentioned above) and thus to better segregate the forest stands close to each other.

As our main purpose was to analyze macro-climatic drivers over large gradients, we assumed that the variations of forest dynamics were more determined by climate average conditions over time than by extreme climatic events, similarly to many species distribution models (Guisan and Thuiller 2005). After a brief analysis of the climate data, we also considered that climate conditions remained stable over time (in particular, we assumed that climate change was negligible over the 20th century), so that we could compare our forest stands despite the differences in terms of time periods (period length, inventory dates). We checked that the length of data periods and the mean of inventory dates were not correlated with calibrated Samsara2 parameters (correlations were lower than 0.3 and not significant in all cases, see Table D.1 in Appendix D)

Finally, two other environmental variables were considered and computed at the stand scale: (1) temperature seasonality, defined as the standard deviation of monthly temperature over year, an indicator of intra-annual variability of temperature (and thus of thermal continentality) that can have important effects on forest composition (Gauquelin and Courbaud 2006; Elkin et al. 2013); and (2) soil pH (pH), which is an important ecological factor for the study species (Piedallu et al. 2009). Soil pH data for French stands were provided by the LerFOB laboratory (Coudun et al. 2006), where it was computed from the French Forest national inventory data by bio-indication. For the Slovenian stands, soil pH was provided with the historical data. The soil pH data were not available for the Swiss stands.

### 4.3 First step: defining and estimating the demographic variables

#### 4.3.1 Definition of empirical and model-based approaches

We quantified the demographic processes from our data using two different approaches. The first method, thereafter named “empirical approach”, consisted in estimating directly stand level demographic fluxes from our data, using population statistics. In the second one, thereafter named “modelbased approach”, we first calibrated the most influent parameters of our individual-based model. We then used these calibrated parameters as demographic variables to analyze climate/demography relations. The main objective of this dual analysis was to show that the model-based approach enables to take into account the effect of stand structure (size distribution, tree density) properly and eliminates its potential confounding effect in the analysis of climate effect. This point is of particular interest in our case because stand structure was highly variable among our stands and generally changed importantly over time.

#### 4.3.2 Computation of population statistics (empirical approach)

We first estimated species dynamics using population statistics computed directly from the raw data as:

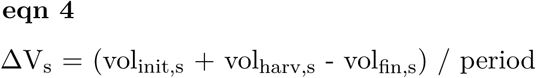
where *ΔV* was an estimate of the annual volume increment; *vol_init_, vol_harv_* and *vo_fin_* were the sum of tree volumes in the oldest inventory, the harvest data and the youngest inventory respectively; *period* was the number of years between the oldest and the youngest inventories, *s* stands for species. Only the trees with DBH above 17.5 cm (20 cm for Slovenian forest) were considered.

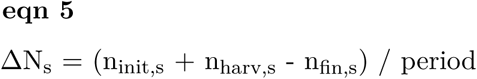
where *ΔN* was an estimate of the species annual recruitment rate, *i.e*. the average number of trees reaching the DBH 17.5 cm (20 cm for Slovenian forest) each year; *n_init_, n_harv_* and *n_fin_* were the number of trees for the species in the oldest inventory, the harvest data and the youngest inventory respectively and *period* was the number of years between the oldest and the youngest inventories, *s* stands for species.

#### 4.3.3 Calibration of the Samsara2 parameters (model-based approach)

Providing detailed information about the Samsara2 model and the calibration step is out of the scope of this study, as it was done elsewhere by Courbaud *et al*. (2015) and Lagarrigues *et al*. (2015) respectively. We nevertheless give here the key points about the Samsara2 model and the calibration procedure. Samsara2 is an individual-based and spatially-explicit model developed to simulate mountain forest dynamics (Courbaud et al. 2015). The demographic processes (tree growth, natural regeneration by germination and tree mortality) and the allometric equations used to deduce the key tree dimensions from DBH are modulated by species-specific parameters (Vieilledent et al. 2010). The initial objective of Samsara2 was to illuminate how forest stand dynamic patterns emerge from the development of individual trees in competition for light, and in return, how the development of individual trees is controlled by stand structure manipulated by forest management. No other environmental drivers than light (and in particular climate) are currently included explicitly in the model.

Here are the main equations for the individual tree growth and regeneration processes, which are of particular interest in this study:

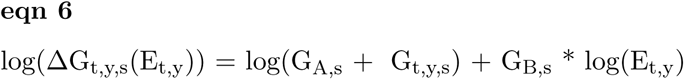
where *ΔG* is the tree basal area annual increment, *G_A_* and *G_B_* are species-specific model parameters, *E* is the amount of light energy intercepted by the tree crown during the annual vegetation period and *G* is a random factor with a zero mean and a species-specific variance, which varies with tree and year to account for individual and annual variabilities of tree growth. The *t, y* and *s* subscripts mean *tree, year* and *species* respectively.

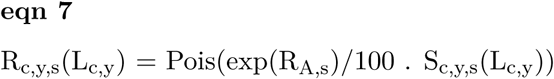
where *R* is the annual regeneration (that is, the number of new saplings recruited in each 10 m × 10 m ground cell), *R_A_* is the log-transformed potential number of saplings (species-specific model parameter) and *S* is the survival rate, *i.e*. a number between 0 and 1 depending on species parameters and on the proportion of light reaching the ground cell during the considered year (L). The factor 100 is a scaled factor, from stand (10,000 m^2^) to ground cell (100 m^2^). The *S* function is modeled by a bell-shaped response to light and is equal to 1 for an intermediate level of light, which differs between species. *Pois* stands for a random draw from a Poisson distribution.

Following the procedure detailed in Lagarrigues et al. (2015), we calibrated for each forest stand two parameters per species, *G_A_* and *R_A_*, using an inverse modeling technique called Approximate Bayesian Computation (ABC). We used rather uninformative priors and ran 10,000 simulations for each stand. We thus performed 1,630,000 simulations in total (10,000 simulations × 163 stands). Each simulation consisted in reproducing the stand historical dynamics and management, starting from the oldest inventory, and then by simulating successively natural dynamics and harvests as monitored in forestry data up to the date of the youngest inventory. The youngest inventory data was used as observational data to estimate the joint posterior distributions of the species model parameters for each stand independently. The modes of posterior distributions were then used as observational data for the following analysis (see the next section). The slight modifications made in the method defined by Lagarrigues et al. (2015) in the course of this study are detailed in the Appendix C.

The individual approach and the detailed modeling of light distribution among the different canopy layers used in the model Samsara2 enabled an explicit representation of the effect of stand structure on the light resource. The stand structure thus affected tree growth and regeneration indirectly by modulating light conditions. For instance, the denser the stand, the less light is available for each tree (E_t,y_ in Eq.6) and the less light reaches the ground and is available for seedlings (L_c,y_ in Eq.7). This main effect of stand structure being taken into account explicitly, we thus expected that the variations of the estimates of the parameters (*G_A_* and *R_A_*) in different stands were little influenced by stand structure and were mainly influenced by other environmental factors.

Concerning the regeneration process, the R_A_ parameter represented the number of saplings potentially recruited as adult trees (*i.e*. once they reach the DBH 7.5 cm) at optimal light conditions after initial reduction due to ground light conditions. Seedling growth spanned over several years in the model before recruitment occurred but seedling mortality caused by non optimal ground light conditions was collapsed into a single event at the same time than germination (Eq. 7). The parameter R_A_ represented thus more the renewal of adult trees (*i.e*. the number of saplings that will reach the adult stage) than a realistic number of saplings. This relative simple representation of the regeneration process was necessary in this study since our data give only information about adult trees and a more sophisticated representation of sapling dynamics can hardly be calibrated with such data.

Regarding the relations to environmental factors, as the R_A_ parameter synthesizes the potential recruitment over long periods (from installation to recruitment), we made the assumption that it would be more sensitive to average climate conditions than to extreme events.

### 4.4 Second step: quantifying the relations between the demographic and the macro-environmental variables

#### 4.4.1 Site definition

From this step on, the stands were grouped in *sites*. The motivation for this aggregation was that some forest inventories were based on statistical sampling, with few points at the stand scale (typically, 10 samples in a stand of 10 ha), making the estimates of the demographic variables more robust statistically at the site scale. Basically, the sites corresponded to forests, except for the large forests encompassing a large altitudinal gradient, which were further divided into multiple sites in order to have homogeneous climatic conditions within sites. This site definition was performed using formal clustering techniques and qualitative considerations (detailed in Appendix A). 31 sites were accordingly defined, they are listed in Table 1 and the stand characteristics are detailed in the Table A.1 (Appendix A). The site-level population statistics and the environmental variables were computed by averaging the stand-level values. The calibrated parameters for a given site were computed by merging the stand-scale posterior distributions and then by taking the mode of the global distribution.

#### 4.4.2 Multiple linear regressions

We carried out regressions between demographic and environmental variables using site-level data. We used multiple linear models, independently for each demographic process (growth and regeneration) and species (spruce, fir and beech), applied first on the population statistics and then on the Samsara2 parameters (*i.e*. twelve independent models in total). We assumed that the site dynamics were independent from each other and we ignored possible spatial autocorrelations. We used the annual sum of growing degree-days (SGDD), the climatic water budget (CWB), the temperature seasonality and the soil pH as explicative variables. We checked that these factors were not correlated with each other (see Appendix D). In addition to these environmental variables, we added an endogenous factor, namely average species basal area, which was computed as the mean of species basal area between the oldest and the youngest inventories.

We added quadratic terms for SGDD and CWB. Quadratic responses of tree growth to these factors have been questioned recently in conditions where thermal or water conditions are limiting (Bugmann and Solomon 2000). However, as we did not study the extreme conditions for our species on these two environmental axes, we considered that the left hand side of a second-degree polynomial response curve was a good approximation.

We applied a procedure of model selection based on the AIC criterion and included only the parsimonious set of significant variables in the final results.

## 5 Results

### 5.1 Variability of the demographic variables

#### 5.1.1 Population statistics

The annual volume increments computed directly from the forestry data ranged from about 3 m^3^.ha^-1^.year^-1^ (in the beech stands and in some high elevation stands) to more than 10 m^3^.ha^-1^.year^-1^ in some very productive spruce-fir stands (Table 1). The annual recruitment rates ranged from close to 0 to 12 trees.ha^-1^.year^-1^. The negative values of recruitment rates mainly resulted from the high uncertainties introduced by the statistical samplings used to inventory some forests (see section 2.4.1).

#### 5.1.2 Samsara2 parameters

The parameters inferred by ABC allowed the Samsara2 model to reproduce adequately the past dynamics of the forest stands, as the model predictions done with calibrated parameters were generally close to the observed data (see Appendix E, Fig. E.1). The Samsara2 regeneration parameters were well correlated with the corresponding population statistics (annual volume increment and annual recruitment rate), but these correlations were weak for the growth process, in particular for spruce and beech (Table 2). Similarly to the population statistics, the Samsara2 parameters were also very variable among stands and sites. Quantitatively, these ranges of parameter values resulted in a tenfold ratio between the minimum and the maximum tree basal area increment for fir and beech (16 for spruce), and a six-fold ratio between the minimum and the maximum annual regeneration for all species (see the parameter values by species and by stand in Appendix E, Table E.1). In order to compare the inter-site and intra-site variabilities, we selected the two sites for which we had the largest numbers of stands (Prénovel 1 and Jura 1). We observed that the intra-site variability was close to the inter-site variability (Fig. 3). The F-test of equality of variances confirmed this result, revealing that the inter-site variance was significantly greater than the Prénovel and Jura 1 intra-site variabilities only for fir growth and spruce regeneration (stars in Fig. 3).

**Table 2:** Correlations between the Samsara2 parameters and the population statistics computed from raw data. The *Process* column identifies the demographic process: for the growth process, the Samsara2 parameter *G_A_* was compared to the population statistic *ΔV*; for the regeneration process, the Samsara2 parameter *R_A_* was compared to the population statistic *ΔN*. The *Correlation* column is the Spearman correlation coefficient, the number of stars indicates the significance (*** for a p-value lower than 0.001, ** lower than 0.01 and * lower than 0.05).

| Species | Process | Correlation |
| --- | --- | --- |
| Beech | Growth | 0.24 |
| Beech | Regeneration | 0.43* |
| Fir | Growth | 0.59*** |
| Fir | Regeneration | 0.82*** |
| Spruce | Growth | -0.25 |
| Spruce | Regeneration | 0.58*** |

**Figure 3:**
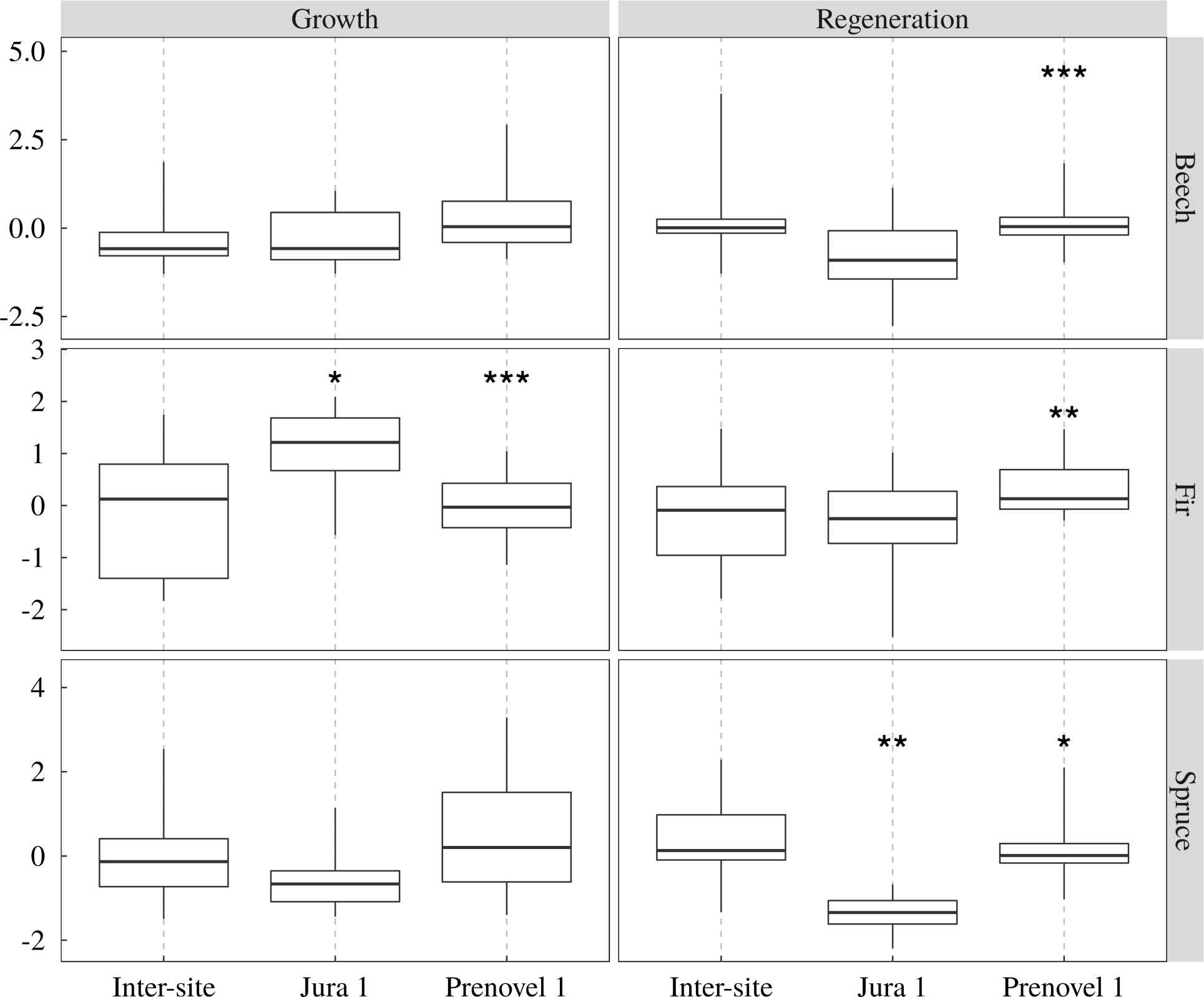
Distribution of growth and regeneration Samsara2 parameters after calibration on forestry data. The *Inter-site* category represents the distribution of the site-level Samsara2 parameters over the 31 sites. The *Jura 1* and *Prénovel 1* categories represent the distribution of Samsara2 parameters for the stands located in the sites Jura 1 and Prénovel 1, respectively. The parameter values (Y axis) were scaled by parameter and species for visualization purpose: V_sc_ = (V - V_m_)/ where *V_sc_* is the scaled value, *V* is the true parameter value, *V_m_* and are respectively the mean and the standard deviation over all stands. The stars above boxplots indicate the significance of the F-test for equality of variances, where we tested if the inter-site variance was larger than the intra-site variance in the “Prénovel 1” and “Jura 1” sites. *** indicates p-value lower than 0.001, ** lower than 0.01 and * lower than 0.05.

### 5.2 Regressions against the environmental variables

#### 5.2.1 Growth

The population statistics describing the growth process (*i.e*. annual volume increment, empirical approach) were mainly, highly and positively related to the average basal area in the site (Table 3). Conversely, the Samsara2 individual-level growth parameters (model-based approach) were not influenced by this factor. Both the fir population statistic and Samsara2 parameter were significantly and positively related to the annual sum of growing degree days (SGDD), but the magnitude of the effect was larger for the Samsara2 parameter (Table 3). We also observed a quadratic but weak relation between the Samsara2 spruce parameter and SGDD (Table 3 and Fig. 4).

**Figure 4:**
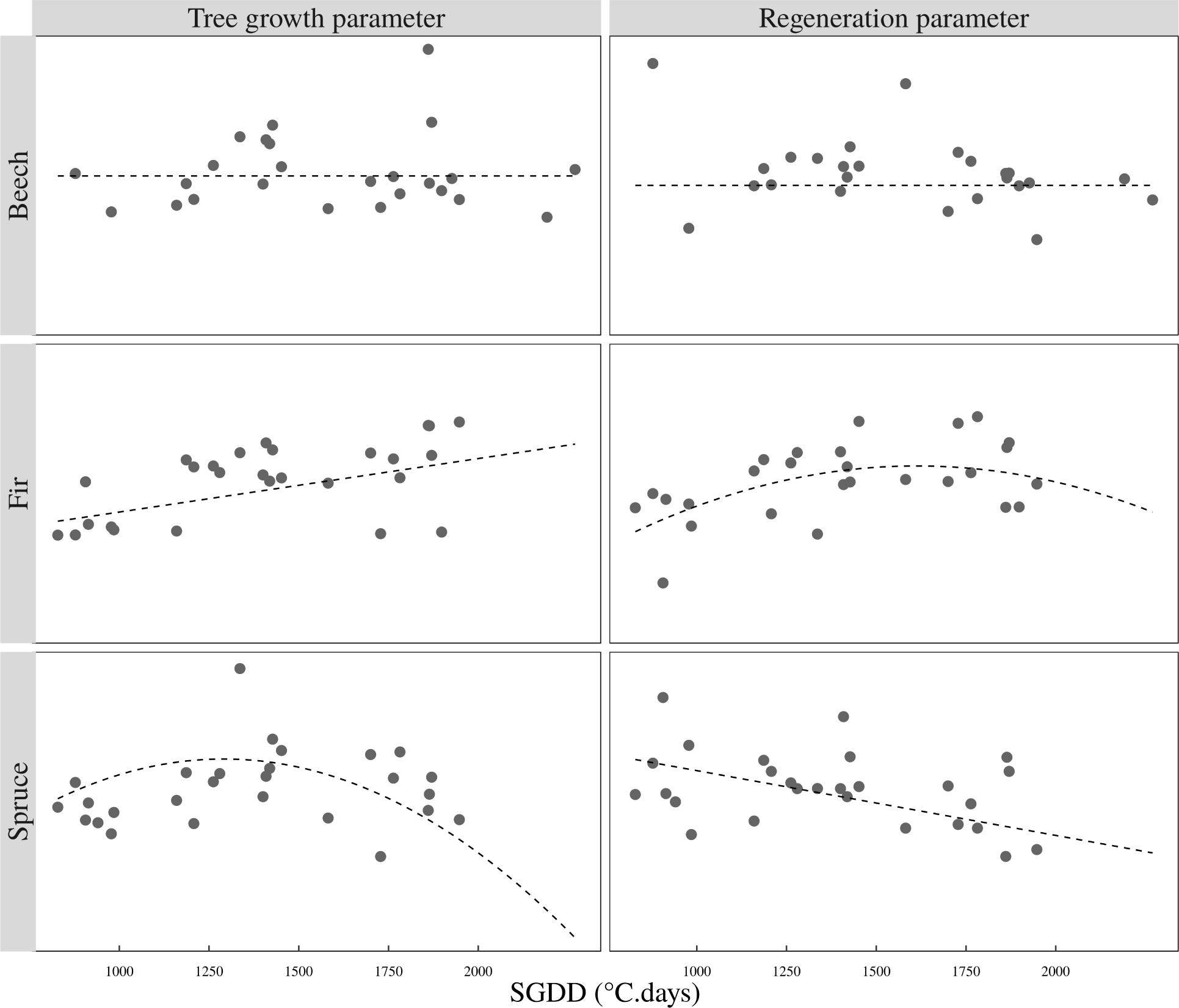
Responses of the site-level Samsara2 parameters to the sum of growing degree-days (SGDD). The dots represent the observed values and the dashed lines represent the responses predicted by the multivariate regression models.

**Table 3 :** Regression results. The values are the magnitudes of effects (*i.e*. the coefficients of the regressions computed on the scaled values), the number of stars indicates the significance of effect (*** for a p-value lower than 0.001, ** lower than 0.01 and * lower than 0.05). The *Process* and *Variable* columns identify the demographic variable used for the regression: *ΔV* and *ΔN* were the population statistics used in the empirical approach (see section 2.3.2); *G_A_* and *R_A_* were the Samsara2 parameters that were calibrated in the model-based approach (see section 2.3.3). BA stands for the average Basal Area in the site; SGDD for the Sum of annual Growing Degree-Days; CWB for the Climatic Water Budget; SEAS for the temperature seasonality; pH for the soil pH. *R^2^* is the adjusted r-squared of regressions. The *n* column indicates the number of sites considered for the regression. In the models where pH was significant, the stands where soil pH was unknown were removed from the observations, which explained, for example, why we had different numbers of points (*n*) between growth and regeneration Samsara2 parameters for fir.

| Process | Species | Variable | Intercept | BA | SGDD | SGDD<sup>2</sup> | CWB | CWB<sup>2</sup> | pH | SEAS | R<sup>2</sup> | n |
| --- | --- | --- | --- | --- | --- | --- | --- | --- | --- | --- | --- | --- |
| Growth | Beech | $\Delta V$ | 0* | 0.9*** | 0 | 0 | 0 | 0 | 0 | 0 | 0.8 | 25 |
| | | $G_A$ | 0 | 0 | 0 | 0 | 0 | 0 | 0.5* | 0 | 0.21 | 18 |
| | Fir | $\Delta V$ | 0 | 0.76*** | 0.24* | 0 | 0 | 0 | 0 | 0 | 0.75 | 26 |
| | | $G_A$ | 0*** | 0 | 0.51*** | 0 | 0 | 0 | 0.36* | 0.57*** | 0.79 | 19 |
| | Spruce | $\Delta V$ | 0 | 1.21*** | 0 | 0 | 0 | 0 | 0.54 | 0 | 0.61 | 17 |
| | | $G_A$ | 0.69* | 0 | -0.32* | -0.73* | 0 | 0 | 0.74*** | 0 | 0.45 | 18 |
| Regeneration | Beech | $\Delta N$ | -0.19* | 0.75*** | -0.39* | 0 | -0.44* | 0.2* | 0.26 | 0 | 0.73 | 18 |
| | | $R_A$ | 0* | 1.18*** | 0 | 0 | 0 | 0 | -0.47* | 1.2*** | 0.41 | 18 |
| | Fir | $\Delta N$ | 0.39* | 0 | 0.39* | -0.4 | 0 | 0 | 0 | 0 | 0.21 | 26 |
| | | $R_A$ | 0.35 | 0 | 0.45* | -0.36 | 0 | 0 | 0 | 0 | 0.27 | 27 |
| | Spruce | $\Delta N$ | -0.39*** | 0 | 0 | 0 | -0.6*** | 0.4*** | 0 | 0 | 0.48 | 26 |
| | | $R_A$ | 0 | 0 | -0.66* | 0 | 0 | 0 | 0.56* | 0 | 0.35 | 18 |

All Samsara2 parameters were positively influenced by soil pH (pH) and the fir Samsara2 parameter was positively related to seasonality (SEAS) (Table 3).

The R^2^ statistics obtained for regressions applied on population statistics were larger than 0.61 and confirmed that the variations of these demographic variables were largely explained by the average basal area (Table 3, white lines within growth section). Interestingly, for the fir growth Samsara2 parameter, we obtained a similar R2 than for the corresponding population statistic, but in that case the main factor was SGDD and not average basal area in the site. The variation of the beech Samsara2 parameter was poorly explained by our set of environmental factors (R^2^ of 0.21).

#### 5.2.2 Regeneration

The regeneration population statistic (*i*.*e*. annual recruitment rate, empirical approach) and Samsara2 regeneration parameter (model-based approach) were strongly and positively related to the average basal area (Table 3) for beech, but not for fir and spruce. The fir regeneration was related to SGDD by a positive saturating response (quadratic response with a maximum obtained for high values of SGDD) (Fig. 4). The spruce regeneration was negatively linked to SGDD. The p-values for these effects were low (Table 3). The other relations between the spruce and beech regeneration population statistics and SGDD or CWB were also weak.

The soil pH and the seasonality had generally no or weak effects on the Samsara2 regeneration parameters.

The R^2^ for the fir regeneration parameter was particularly low, highlighting the low influence of SGDD even if this factor was retained in the model selection (Table 3).

## 6 Discussion

### 6.1 Using the model-based demographic variables enables to factor out the effects of stand density and structure on tree growth

The population annual volume increment (empirical approach) was highly correlated with the population basal area, whereas the tree-level growth parameters (model-based approach) did not depend on this factor (Table 3). The independence between the tree growth parameter *G_A_*, which is the scale parameter in the Samsara2 tree growth equation (Eq. 6), and the population density confirms that the density dependence of population growth is adequately modeled in Samsara2, through the effect of stand structure on the distribution of light among trees. Moreover, the Samsara2 parameters were poorly correlated with the annual volume increment computed from raw data (Table 2). This means that the model-based approach enabled us to factor out the effects of stand density and structure on tree growth, and thus to detect the potential climatic drivers of forest dynamics better than previous studies based on simpler dynamic models (*e.g*. (Thuiller et al. 2014)). We were able to draw such demographic inferences from forestry data thanks to an efficient inverse calibration technique (ABC). Due to its flexibility, ABC can indeed make use of original types of data that do not contain all the necessary ingredients to perform standard inferences. In our case, we could infer one growth and one regeneration parameters by species from forestry data, since we were able to simulate with the Samsara2 model both the biological dynamics of forest trees and the stand specific harvesting treatments imposed by foresters (Lagarrigues et al. 2015).

Concerning regeneration, we did not find significant correlations between population statistics (*i.e*. the annual recruitment rate) and population basal area. This indicates that tree density is not a key driver of recruitment rate at stand scale. Besides, the Samsara2 regeneration parameters (model-based approach) were significantly correlated with the stand level annual recruitment rates (empirical approach) (Table 2). That result either showed that stand structure does not affect significantly the regeneration process either, or that the regeneration process as it was modeled in our model-based approach is too simplistic to take into account size structure properly. Our data did not allow us to disentangle these two competing explanations.

### 6.2 Responses of tree demography to macro-environmental variables

An important result of this study is that the inter-site variability of Samsara2 growth and regeneration parameters (model-based approach) was not significantly larger than the intra-site variability, except for fir growth and spruce regeneration (Fig. 3). A possible explanation of this finding is that micro-environmental conditions can be highly variable within mountain forest stands, because of aspect heterogeneity, altitudinal ranges and the presence of crests and little depressions that create large differences in soil depth and fertility. Such physical heterogeneity is known to cause huge differences in community assembly and functioning for low-stature mountain plants (Scherrer and Körner 2011). It is likely to cause significant differences for trees also, at least during their early stages. The important local drivers of this intra-site variability should be further studied, and the consequences of this variability on forest dynamics should be better taken into account when projecting forest dynamics, notably under climate change scenarios. The inverse calibration approach detailed in this study provides concrete ways to propagate this intra-site variability in dynamical projections.

The responses of Samsara2 growth parameters to thermal conditions (SGDD) were contrasted between species, in terms of shape (flat for beech, linear for fir or quadratic for spruce) (Fig. 4) and of significance (high for fir, weak for spruce) (Table 3). This result contradicts the high and continuous positive response of growth to temperature usually assumed for all species in DVMs in mountain environments (Bugmann and Solomon 2000; Lexer and Honninger 2001; Bugmann 2001; Seidl et al. 2005), which thus projected strong effects of climate change on forest dynamics (Elkin et al. 2013; Seidl and Lexer 2013). Interestingly, SGDD had also contrasted and weak effects on natural regeneration in our study. It is noteworthy that other temperature indicators that are known to impact the demography of these species (such as the winter minimum temperature for tree establishment (Bugmann 1996) or the spring temperature for tree growth (Battipaglia et al. 2009)) were highly correlated with SGDD (0.91 and 0.99 respectively, see Appendix D). This means that the poor predictive power of SGDD can be interpreted as a poor predictive power of temperature in general (except the temperature seasonality, see below). Importantly, this result is not in contradiction with the threshold effect of winter temperature on the occurrence of species establishment usually used in DVMs (Bugmann 2001), since our study focused on non-extreme conditions within the thermal envelope of the species (Fig. 2). Besides, we also obtained a negative relation between spruce regeneration and SGDD (Table 3). This result confirms the preference of spruce for cold environments, but our data did not allow determining if it is due to spruce intrinsic fitness or to the effect of competition with saplings of other species. Finally, our results showed that temperature may not be a key factor to understand species demography in forest stands where temperature is not a strong limiting factor. Other factors may therefore be preponderant for tree community dynamics in these forests, such as management (Carcaillet and Muller 2005) or competition for resources (Kunstler et al. 2011). The interactions between macro-environmental variables and such other factors (*e.g*. (Lévesque et al. 2016)) should therefore be better taken into account in DVMs.

Natural regeneration was found to be little driven by the macro-environmental factors used in this study (Table 3). This finding is consistent with the fact that germination is highly sensitive to the presence of micro-sites (Prach et al. 1996; Hunziker and Brang 2005). Another limiting factor for regeneration is the browsing by large herbivores, particularly for silver fir. This browsing pressure greatly varies between the studied forests: whereas it is not yet considered as limiting in the Western Alps, fir regeneration is already highly impacted in the Central and Eastern Alps (Hunziker and Brang 2005; Diaci et al. 2011). These two factors may have larger effects on tree regeneration than climatic factors (Didion et al. 2011).

As expected, the water conditions had little influence (Table 3), probably because our stands were located in well-watered conditions (Fig. 2). In a context of global change, this should be no longer the case in the future, particularly at low elevations (Wilson and Hopfmueller 2001). Here again, the absence of response may be a sign of a threshold effect: no influence in a large range of conditions where water is not limiting, and rapid decrease where the conditions become harsh. We were not able to test this hypothesis in this study because of the absence of data in water-limiting conditions. Moreover, we were not able to quantify accurately the water conditions experienced by the trees because of a lack of information about soil properties. We had then to use a simple climatic water budget index without taking the role of soil in the water cycle into account (Granier et al. 2007). Yet, soil is known to play an important role in the water cycle of forest ecosystems (Piedallu et al. 2013). Going further in studying the effect of water conditions on forest demography with such data would thus impose both to collect data in water-limiting conditions and to measure key soil properties in all stands.

Seasonality of temperatures and soil pH have already been used to understand species distribution (Gauquelin and Courbaud 2006; Piedallu et al. 2009) but their explicit relations with demography remain, to our knowledge, unknown. In this study, we observed that seasonality and soil pH had positive effects on the Samsara2 parameters in many cases, suggesting an affinity to more continental and more alkaline conditions, respectively. Nevertheless, these correlations should be considered with care because these factors were here used to segregate two groups (approximately, low seasonality in the Pyrenees vs high seasonality in the Alps, low pH in acid siliceous soils vs high pH in neutral calcareous soils), with a very low variability inside these groups as compared to the gaps occurring between groups (Table A.1 in Appendix A and Fig. D.1 in Appendix D). These relations should thus be investigated with a better sample of sites along these factors.

### 6.3 The potential of forestry data for ecological research

This study relied on data mainly collected by forest managers to monitor the evolution of their forests rather than on research data. Considering the data scarcity in forest ecology, these forestry data are useful to quantify forest dynamics along large environmental gradients. In particular, an important added value of these long term data (up to 100 years of forest dynamics) is to allow the quantification of the renewal of forest stands by means of natural regeneration. Indeed, the lack of long-term data is often an important limit to predict the future species dynamics in forest stands (Piedallu et al. 2009; Snell et al. 2014). Besides, this data source is far from being completely exploited with this study: further information can likely be extracted from the present data set (*e.g*. to study the mortality process (Csillery et al. 2013)) and management documents are available for many other forests across Europe. Thanks to modern inference techniques such as ABC, forestry data give critical information about the long-term forest dynamics in a warming climate, and should therefore be more widely compiled and used for global change research.

## 7 Acknowledgment

We thank all data providers, in particular the French Forest National Office for its assistance while extracting data from their archives. The data compilation for the Slovenian stands, done by the University of Ljubljana, was funded by the European FP7 research project “Advanced multifunctional forest management in European mountain ranges” (ARANGE, n° 289437). The simulations were carried out using the computing center CIMENT located in the Grenoble University and we thank the CIMENT supporting team and Eric Maldonado from IRSTEA Grenoble for their help to fix technical issues related to the use of the simulation platform. Guillaume Lagarrigues was funded by the French Environment and Energy Management Agency (ADEME), the French Forest National Office (ONF) and IRSTEA. IRSTEA Grenoble is a part of labex OSUG@2020. We have no conflicts of interest to disclose.

## 8 Data accessibility

- Forestry and environmental data: will be stored in Dryad
- Samsara2 model source code: available in the Capsis platform (Dufour-Kowalski et al. 2012) (http://capsis.cirad.fr/), with restricted access
- R scripts: uploaded as online supporting information

## 9 Supporting information

Additional supporting information may be found in the online version of this article:

- Appendix A: Stand and site features
- Appendix B: Temperature correction in function of elevation
- Appendix C: Specific recalibration features for this study
- Appendix D: Correlations between environmental variables
- Appendix E: Detailed results of the recalibration step

## References

Allen RG, Smith M, Raes D, Pereira LS (1998) Crop evapotranspiration: guidelines for computing crop water requirements. Food; Agriculture Organization of the United Nations, Rome

Battipaglia G, Saurer M, Cherubini P et al (2009) Tree rings indicate different drought resistance of a native (Abies alba Mill.) and a nonnative (Picea abies (L.) Karst.) species co-occurring at a dry site in Southern Italy. Forest Ecology and Management 257:820–828. doi: 10.1016/j.foreco.2008.10.015

Bodin J, Badeau V, Bruno E et al (2013) Shifts of forest species along an elevational gradient in Southeast France: climate change or stand maturation? Journal of Vegetation Science 24:269–283. doi: 10.1111/j.1654-1103.2012.01456.x

Bugmann H (2001) A review of forest gap models. Climatic Change 51:259–305. doi: 10.1023/A:1012525626267

Bugmann HK (1996) A simplified forest model to study species composition along climate gradients. Ecology 77:2055–2074.

Bugmann HKM, Solomon AM (2000) Explaining Forest Composition and Biomass across Multiple Biogeographical Regions. Ecological Applications 10:95–114. doi: 10.2307/2640989

Büntgen U, Frank DC, Kaczka RJ et al (2007) Growth responses to climate in a multi-species tree-ring network in the Western Carpathian Tatra Mountains, Poland and Slovakia. Tree Physiology 27:689–702.

Carcaillet C, Muller SD (2005) Holocene tree-limit and distribution of Abies alba in the inner French Alps: anthropogenic or climatic changes? Boreas 34:468–476.

Carrer M, Motta R, Nola P (2012) Significant Mean and Extreme Climate Sensitivity of Norway Spruce and Silver Fir at Mid-Elevation Mesic Sites in the Alps. PLoS one. doi: 10.1371/journal.pone.0050755

Coudun C, Gégout J-C, Piedallu C, Rameau J-C (2006) Soil nutritional factors improve models of plant species distribution: an illustration with Acer campestre (L.) in France. Journal of Biogeography 33:1750–1763. doi: 10.1111/j.1365-2699.2005.01443.x

Courbaud B, Lafond V, Lagarrigues G et al (2015) Applying ecological model evaludation: Lessons learned with the forest dynamics model Samsara2. Ecological Modelling 314:1–14.

Csillery K, Seignobosc M, Lafond V et al (2013) Estimating long-term tree mortality rate time series by combining data from periodic inventories and harvest reports in a Bayesian state-space model. Forest Ecology and Management 292:64–74. doi: 10.1016/j.foreco.2012.12.022

Diaci J, Rozenbergar D, Anic I et al (2011) Structural dynamics and synchronous silver fir decline in mixed old-growth mountain forests in Eastern and Southeastern Europe. Forestry 84:479–491. doi: 10.1093/forestry/cpr030

Didion M, Kupferschmid AD, Lexer MJ et al (2009) Potentials and limitations of using large-scale forest inventory data for evaluating forest succession models. Ecological Modelling 220:133–147. doi: 10.1016/j.ecolmodel.2008.09.021

Didion M, Kupferschmid AD, Wolf A, Bugmann H (2011) Ungulate herbivory modifies the effects of climate change on mountain forests. Climatic Change 109:647–669. doi: 10.1007/s10584-011-0054-4

Dittmar C, Zech W, Elling W (2003) Growth variations of Common beech (Fagus sylvatica L.) under different climatic and environmental conditions in Europe—a dendroecological study. Forest Ecology and Management 63–78.

Dufour-Kowalski S, Courbaud B, Dreyfus P et al (2012) Capsis: an open software framework and community for forest growth modelling. Annals of Forest Science 69:221–233.

Durand Y, Brun E, Mérindol L et al (1993) A meteorological estimation of relevant parameters for snow models. Annals of Glaciology 18:65–71.

Elkin C, Giuggiola A, Rigling A, Bugmann H (2015) Short- and long-term efficacy of forest thinning to mitigate drought impacts in mountain forests in the European Alps. Ecological Applications 25:1083–1098. doi: 10.1890/14-0690.1

Elkin C, Gutiérrez AG, Leuzinger S et al (2013) A 2 °C warmer world is not safe for ecosystem services in the European Alps. Global Change Biology 19:1827–1840. doi: 10.1111/gcb.12156

Fontes L, Bontemps JD, Bugmann H et al (2010) Models for supporting forest management in a changing environment. Forest Systems 19:8–29.

Gauquelin X, Courbaud B (2006) Guide des sylvicultures de montagne. ONF/CRPF/Cemagref

Granier A, Reichstein M, Bréda N et al (2007) Evidence for soil water control on carbon and water dynamics in European forests during the extremely dry year: 2003. Agricultural and Forest Meteorology 143:123–145. doi: 10.1016/j.agrformet.2006.12.004

Grassi G, Minotta G, Tonon G, Bagnaresi U (2004) Dynamics of Norway spruce and silver fir natural regeneration in a mixed stand under uneven-aged management. Canadian Journal of Forest Research-Revue Canadienne De Recherche Forestiere 34:141–149. doi: 10.1139/x03-197

Guisan A, Thuiller W (2005) Predicting species distribution: offering more than simple habitat models. Ecology Letters 8:993–1009. doi: 10.1111/j.1461-0248.2005.00792.x

Hanewinkel M, Cullmann DA, Schelhaas M-J et al (2013) Climate change may cause severe loss in the economic value of European forest land. Nature Climate Change 3:203–207. doi: 10.1038/nclimate1687

Hartig F, Dyke J, Hickler T et al (2012) Connecting dynamic vegetation models to data - an inverse perspective. Journal of Biogeography 39:2240–2252. doi: 10.1111/j.1365-2699.2012.02745.x

Hickler T, Vohland K, Feehan J et al (2012) Projecting the future distribution of European potential natural vegetation zones with a generalized, tree species-based dynamic vegetation model. Global Ecology and Biogeography 21:50–63. doi: 10.1111/j.1466-8238.2010.00613.x

Hijmans RJ, Cameron SE, Parra JL et al (2005) Very high resolution interpolated climate surfaces for global land areas. International journal of climatology 25:1965–1978.

Hunziker U, Brang P (2005) Microsite patterns of conifer seedling establishment and growth in a mixed stand in the southern Alps. Forest Ecology and Management 210:67–79.

Keenan RJ (2015) Climate change impacts and adaptation in forest management: a review. Annals of Forest Science 72:145–167.

Klopcic M, Jerina K, Boncina A (2010) Long-term changes of structure and tree species composition in Dinaric uneven-aged forests: are red deer an important factor? European Journal of Forest Research 129:277–288. doi: 10.1007/s10342-009-0325-z

Körner C, Basler D, Hoch G et al (2016) Where, why and how? Explaining the low-temperature range limits of temperate tree species. Journal of Ecology 104:1076–1088. doi: 10.1111/1365-2745.12574

Kunstler G, Albert CH, Courbaud B et al (2011) Effects of competition on tree radial-growth vary in importance but not in intensity along climatic gradients. Journal of Ecology 99:300–312.

Lagarrigues G, Jabot F, Lafond V, Courbaud B (2015) Approximate Bayesian computation to recalibrate individual-based models with population data: Illustration with a forest simulation model. Ecological Modelling 306:278–286.

Lenoir J, Gégout J-C, Pierrat J-C et al (2009) Differences between tree species seedling and adult altitudinal distribution in mountain forests during the recent warm period (1986-2006). Ecography 32:765–777.

Lexer MJ, Honninger K (2001) A modified 3D-patch model for spatially explicit simulation of vegetation composition in heterogeneous landscapes. Forest Ecology and Management 144:43–65. doi: 10.1016/s0378-1127(00)00386-8

Lévesque M, Walthert L, Weber P (2016) Soil nutrients influence growth response of temperate tree species to drought. Journal of Ecology 104:377–387. doi: 10.1111/1365-2745.12519

Lindner M, Fitzgerald JB, Zimmermann NE et al (2014) Climate change and European forests: What do we know, what are the uncertainties, and what are the implications for forest management? Journal of environmental management 146:69–83.

Mäkinen H, Nöjd P, Kahle HP et al (2002) Radial growth variation of Norway spruce (Picea abies (L.) Karst.) across latitudinal and altitudinal gradients in central and northern Europe. Forest Ecology and Management 171:243–259.

Mencuccini M, Piussi P, Zanzi Sulli A (1995) Thirty years of seed production in a subalpine Norway spruce forest: patterns of temporal and spatial variation. Forest Ecology and Management 76:109–125. doi: 10.1016/0378-1127(95)03555-O

Oudin L, Hervieu F, Michel C et al (2005) Which potential evapotranspiration input for a lumped rainfall–runoff model?: Part 2—Towards a simple and efficient potential evapotranspiration model for rainfall–runoff modelling. Journal of Hydrology 303:290–306.

Paluch JG, Jastrzębski R (2013) Natural regeneration of shade-tolerant Abies alba Mill. in gradients of stand species compositions: Limitation by seed availability or safe microsites? Forest Ecology and Management 307:322–332.

Penman HL (1948) Natural Evaporation from Open Water, Bare Soil and Grass. Proceedings of the Royal Society of London A: Mathematical, Physical and Engineering Sciences 193:120–145. doi: 10.1098/rspa.1948.0037

Piedallu C, Gégout J-C, Perez V, Lebourgeois F (2013) Soil water balance performs better than climatic water variables in tree species distribution modelling. Global Ecology and Biogeography 22:470–482.

Piedallu C, Perez V, Gégout J-C et al (2009) Impact potentiel du changement climatique sur la distribution de l’Epicéa, du Sapin, du Hêtre et du Chêne sessile en France. Revue forestière française 61:567–593.

Pinto PE, Gégout J-C, Hervé J-C, Dhôte J-F (2008) Respective importance of ecological conditions and stand composition on Abies alba Mill. dominant height growth. Forest Ecology and Management 255:619–629. doi: 10.1016/j.foreco.2007.09.031

Prach K, Lepš J, Michálek J (1996) Establishment of Picea abies Seedlings in a Central European Mountain Grassland: An Experimental Study. Journal of Vegetation Science 7:681–684. doi: 10.2307/3236379

Rolland C, Petitcolas V, Michalet R (1998) Changes in radial tree growth for Picea abies, Larix decidua, Pinus cembra and Pinus uncinata near the alpine timberline since 1750. Trees 13:40–53.

Scheiter S, Langan L, Higgins SI (2013) Next-generation dynamic global vegetation models: learning from community ecology. New Phytologist 198:957–969. doi: 10.1111/nph.12210

Scherrer D, Körner C (2011) Topographically controlled thermal-habitat differentiation buffers alpine plant diversity against climate warming. Journal of Biogeography 38:406–416. doi: 10.1111/j.1365-2699.2010.02407.x

Seidl R, Lexer MJ (2013) Forest management under climatic and social uncertainty: Trade-offs between reducing climate change impacts and fostering adaptive capacity. Journal of Environmental Management 114:461–469.

Seidl R, Lexer MJ, Jäger D, Hönninger K (2005) Evaluating the accuracy and generality of a hybrid patch model. Tree Physiology 25:939–951. doi: 10.1093/treephys/25.7.939

Selås V, Piovesan G, Adams JM, Bernabei M (2002) Climatic factors controlling reproduction and growth of Norway spruce in southern Norway. Canadian Journal of Forest Research 32:217–225. doi: 10.1139/x01-192

Seynave I, Gegout J-C, Herve J-C, Dhote J-F (2008) Is the spatial distribution of European beech (Fagus sylvatica L.) limited by its potential height growth? Journal of Biogeography 35:1851–1862. doi: 10.1111/j.1365-2699.2008.01930.x

Snell R, Huth A, Nabel J et al (2014) Using dynamic vegetation models to simulate plant range shifts. Ecography 37:1184–1197.

Stancioiu PT, O’Hara KL (2006) Regeneration growth in different light environments of mixed species, multiaged, mountainous forests of Romania. European Journal of Forest Research 125:151–162. doi: 10.1007/s10342-005-0069-3

Tegel W, Seim A, Hakelberg D et al (2014) A recent growth increase of European beech (Fagus sylvatica L.) at its Mediterranean distribution limit contradicts drought stress. European Journal of Forest Research 133:61–71.

Thuiller W, Münkemüller T, Schiffers KH et al (2014) Does probability of occurrence relate to population dynamics? Ecography 37:1155–1166.

Tinner W, Colombaroli D, Heiri O et al (2013) The past ecology of Abies alba provides new perspectives on future responses of silver fir forests to global warming. Ecological Monographs 83:419–439.

Vieilledent G, Courbaud B, Kunstler G et al (2010) Individual variability in tree allometry determines light resource allocation in forest ecosystems: a hierarchical Bayesian approach. Oecologia 163:759–773. doi: 10.1007/s00442-010-1581-9

Wang WJ, He HS, Spetich MA et al (2013) A large-scale forest landscape model incorporating multi-scale processes and utilizing forest inventory data. Ecosphere. doi: 10.1890/es13-00040.1

Wilson RJ, Hopfmueller M (2001) Dendrochronological investigations of Norway spruce along an elevational transect in the Bavarian Forest, Germany. Dendrochronologia 19:67–79.

Wisz MS, Pottier J, Kissling WD et al (2013) The role of biotic interactions in shaping distributions and realised assemblages of species: implications for species distribution modelling. Biological Reviews of the Cambridge Philosophical Society 88:15–30. doi: 10.1111/j.1469-185X.2012.00235.x

Zingg A, Frutig F, Bürgi A et al (2009) Yield performance in the plenter forest research plots in Switzerland. Dauerwald 160:162–174. doi: 10.3188/szf.2009.0162

